# Optimization of protocols for immunohistochemical assessment of enteric nervous system in formalin fixed human tissue

**DOI:** 10.1101/2024.12.15.628584

**Authors:** Su Min Hong, Xia Qian, Vikram Deshpande, Subhash Kulkarni

**Author notes:** Address all correspondence to: Subhash Kulkarni, Ph.D.

## Abstract

Gastrointestinal (GI) motility is regulated in a large part by the cells of the enteric nervous system (ENS), suggesting that ENS dysfunctions either associate with, or drive GI dysmotility in patients. However, except for select diseases such as Hirschsprung’s Disease or Achalasia that show a significant loss of all neurons or a subset of neurons, our understanding of human ENS histopathology is extremely limited. Recent endoscopic advances allow biopsying patient’s full thickness gut tissues, which makes capturing ENS tissues simpler than biopsying other neuronal tissues, such as the brain. Yet, our understanding of ENS aberrations observed in GI dysmotility patients lags behind our understanding of central nervous system aberrations observed in patients with neurological disease. Paucity of optimized methods for histopathological assessment of ENS in pathological specimens represent an important bottleneck in ascertaining how the ENS is altered in diverse GI dysmotility conditions. While recent studies have interrogated ENS structure in surgically resected whole mount human gut, most pathological specimens are banked as formalin fixed paraffin embedded (FFPE) tissue blocks – suggesting that methods to interrogate ENS in FFPE tissue blocks would provide the biggest impetus for ENS histopathology in a clinical setting. In this report, we present optimized methods for immunohistochemical interrogation of the human ENS tissue on the basis of >25 important protein markers that include proteins expressed by all neurons, subset of neurons, hormones, and neurotransmitter receptors. This report provides a resource which will help pathologists and investigators assess ENS aberrations in patients with various GI dysmotility conditions.

## Introduction

Gastrointestinal (GI) dysmotility disorders afflict large numbers of patients, whose persistent GI dysmotility causes a loss of quality of life and confers them with a significant economic burden^1,2^. GI motility, amongst other gut functions such as immunity, secretion, absorption and interactions with microbiota, is regulated by the actions of the neurons and glial cells of the enteric nervous system (ENS) that is organized in networks called plexuses which are located entirely within the gut wall^3–11^. The near-comprehensive regulation of GI functions, especially its motility, by ENS suggests that the ENS in patients with GI dysmotility disorders would be significantly negatively impacted.

ENS aberrations are observed in patients with congenital Hirschsprung’s Disease, where a developmental defect leads to incomplete innervation of the distal colon leading to a lack of all ENS in the distal colon resulting in severe dysmotility in newborns^12,13^. Aberrant ENS innervation is also observed in patients suffering from achalasia, where a significant loss of inhibitory NOS1 (nitric oxide synthase 1)-expressing nitrergic neurons (those releasing nitric oxide (NO) as a principal neurotransmitter) in the lower esophageal sphincter is associated with impaired esophageal dysmotility^14,15^. However, while GI dysmotility occurs in many other diseases, including in rheumatological diseases such as systemic sclerosis, neurological diseases such as Parkinson’s’ Disease, and GI diseases and syndromes such as inflammatory bowel disease (IBD), idiopathic and diabetic gastroparesis, and irritable bowel syndrome (IBS), the nature of the ENS aberrations in these patients remains unknown^1,16–18^. The lack of data about the nature of ENS defects in these diseases does not necessarily signify a normal ENS in these patients, but instead suggests that efforts are needed to better interrogate the tissues from patients with and without GI dysmotility to catalogue if the ENS tissue in dysmotility patients suffers from a structural loss of neuronal numbers or nerve fibers, or whether the pathology results from altered expression of important neurotransmitters, receptors, hormones, and other markers.

An important bottleneck in characterizing the ENS defects in GI dysmotility patients has been a lack of optimized histochemical protocols for characterizing ENS tissue from patients. Formalin fixed paraffin embedded (FFPE) tissue blocks are the mainstay of archival tissues used by pathological labs and medical centers, and a study at the turn of the century estimated that biobanks contain more than 300 million tissue blocks, with their numbers growing by 20 million a year^19^. Recently, it has been reported that these numbers are almost certainly an underestimation, and that archived human tissue in FFPE blocks allow for high quality preservation of cellular structure and proteomic and transcriptomic data at ambient temperatures^20,21^. Biorepositories containing tissue specimens in FFPE blocks from well-phenotyped patients thus provides an invaluable resource to interrogate the ENS in dysmotility patients using histopathological methods. Thus, it is imperative that robust, detailed, and standardized immunohistochemical protocols be developed for assessing the expression of important ENS and other pathological markers in the human ENS tissue. To this, this study presents detailed protocols for immunostaining human ENS tissues obtained from archived FFPE tissue blocks with antibodies against >25 different neuronal, glial, and other pathological markers that are relevant to ENS structure, function, and may be implicated in disease.

## Materials

### Human tissue procurement and immunohistochemistry

Paraffin sections of the adult small intestinal full-thickness gut from deidentified human tissues were obtained from the Department of Pathology at Beth Israel Deaconess Medical Center, and the Institutional Review Board approved using the deidentified human tissues.

Since we wish to control for the influence of materials on the immunohistochemical staining of tissue sections, we list in detail all the materials used in this study.

1. Microtome

a. Shandon Cat. No. AS325
2. Lighted tissue bath

a. Premiere Lighted Tissue Bath Cat. No. XH 1003
3. <u>Staining slides</u>: Colorfrost microscope slides (Fisher Scientific; Cat. No.1255018)
4. <u>Slide staining chambers</u>: EasyDip™ Slide staining jars (Simport; Cat. No. M900-12P)
5. Deparaffinization/Rehydration chemicals

a. Xylene ACS: (Pharmco-Greenfield; Cat. No. 399000000CSFF)
b. Ethanol 200 proof ACS USP: (Pharmco-Greenfield; Cat. No. 111000000CSGL)
6. Heat-mediated antigen retrieval buffer and hardware

a. Citrate Buffer pH6:

i. Sodium citrate: (JT Baker; Cat. No. 2646-01)
ii. Citrate acid: (Fisher Scientific; Cat. No. BP39-500)
b. Tris-EDTA Buffer pH9:

i. Tris: (EMD Millipore; Cat. No. 648310-500GM)
ii. EDTA: (Sigma Aldrich; Cat. No. 56758-100G)
c. Pressure cooker

i. Biocare medical, antigen retrieval chamber (Cat. No. DCARC0001)
7. Blocking Buffer:

a. Normal Horse Serum Control: (Invitrogen; Cat. No. 31874)
b. TritonTM X-100: (Sigma; Cat. No. T8787-50ML)
8. Hydrogen peroxide blocking buffer

a. Hydrogen peroxide 30% (Fisher Bioreagent; Cat. No. 7722-84-1)
9. Secondary antibody:

a. Goat Biotinylated anti-rabbit antibody: (Vector Labs; Cat. No. BA-1000)
b. Horse Biotinylated anti-mouse antibody: (Vector Labs; Cat. No. BA-2001)
10. Avidin-Biotin Complex:

a. ABC kit, (Vector Labs; Cat. No. PK-6100)
11. DAB kit

a. DAB kit: (Vector Labs; Cat. No. SK-4100)
12. Hematoxylin counterstain

a. Gill’s hematoxylin solution No.2. (Electron microscopy science; Cat. No. 26030-24)
13. Mounting media

a. Cytoseal 60 (Epredia; Cat. No. 8310-4)
b. Microscope cover glass; (Alkali scientific Inc; Cat. No. SM2460)
c. Prolong Gold Antifade Reagent: (Invitrogen; Cat. No. P36930)

## Methods

1. After obtaining FFPE tissue blocks for sectioning, the blocks need to be prepared for sectioning. For this, begin by softening the paraffin block. Place the FFPE block in icy water till soft. Depending on the block, this process may take anywhere between 15 minutes to 2 hours. Once the block has softened, use the microtome to begin sectioning. Sectioning is performed to achieve 5 μm thickness of the sections. Carefully spread the sections onto a prewarmed water bath set to 38°C. Once the sections are nicely spread, transfer them onto histology slides. Allow the slides to dry in the dark for two days before staining.
Note: Please consult your histologist for technical expertise in sectioning and placing the sections on the slides.
2. To process the slides, initiate by drying them at 37°C in an oven overnight. Following this, perform a baking step at 60°C for 10 minutes.
3. Proceed with the deparaffinization of FFPE slides using the following step-wise protocol:

a. Two Xylene washes for the slide, each wash is for the duration of 5 minutes.
b. Two 100% Ethanol washes for the slide, each wash is for the duration of 5 minutes.
c. Two 95% Ethanol washes for the slide, each wash is for the duration of 5 minutes.
d. Two 70% Ethanol washes for the slide, each wash is for the duration of 5 minutes.
e. Two double distilled water (ddH2O) washes for the slide, each wash is for the duration of 5 minutes.
4. Next, perform heat-mediated antigen retrieval by placing the slides into a preheated (68°C) pressure cooker, and run the pressure cooker for the designated amount of time and temperature as listed in Table 1.
<u>Note</u>: This step varies with the antibody used to detect the specific antigen. Please check Table 1 for the details on which antigen-retrieval buffer, temperature, and the treatment time is applicable for the antigen to be detected.
5. Allow the slides to cool down in an icy water bath containing slushy ice water for 10 minutes. It is imperative to refrain from removing the slides while the buffer remains hot, as elevated temperatures may cause the tissues to detach from the slides. Finally, wash the slides with 1x phosphate-buffered saline (PBS) once.
Note: Do not use glass slide chambers for this cool-down in slushy ice water.
6. To prepare for blocking, start by making a blocking solution that consists of 5% normal horse serum (NHS), 1X PBS, and 0.5% Triton X-100. Next, use a hydrophobic pen to encircle the tissue on the slide. Carefully add 100 µl of the blocking solution to the slides dropwise, taking care not to pipette it directly onto the tissue, as this can cause the tissue to lift off the slide. Allow the blocking solution to incubate for 1 hour at room temperature. After the incubation period, gently remove the blocking buffer from the slides.
7. Dilute the primary antibody in 1X PBS, then gently add 100 µl of the antibody solution in a dropwise manner. Incubate the slides in a humid chamber, in the dark, at 4°C overnight.
8. Carefully decant the primary antibody solution from each slide the next morning and wash them once with 1X PBS for 5 minutes. After that, proceed with the peroxidase-blocking step. Dilute 30% hydrogen peroxide to a ratio of 1:100 and block the slides for 12 minutes. After the blocking, wash away any excess hydrogen peroxide by rinsing the slides with 1X PBS twice, with each wash lasting 5 minutes.
9. Next, dilute the secondary antibody in 1X PBS, then gently add 100ul of the antibody solution in a dropwise manner. Incubate the slides in a humid chamber, in the dark, at room temperature for 1 hour.
10. Gently decant the secondary antibody solution from each slide and wash them three times with 1X PBS for 5 minutes each. After washing, proceed to the Avidin-Biotin Complex (ABC) step. Prepare the ABC solution 30 minutes before use. Add 100 µl of the ABC solution to the slides in a dropwise manner. Incubate the slides in a humid chamber, in the dark, at room temperature for 30 minutes.
11. Carefully decant the ABC solution from each slide and wash them 3 times with 1X PBS for 5 minutes.
12. After washing, proceed to the DAB chromogen staining step. Prepare the DAB solution and apply it to the tissue section. Assess the intensity of the chromogen staining under the microscope. The staining process may take up to 10 minutes. Once the staining has reached its optimal level, place the slides in a chamber with tap water to stop the reaction. Wash the slides with cold tap water at least 5 times to fully remove residual DAB.
<u>Note</u>: Each antibody will have a different exposure time. For the ENS-specific antibody, please refer to Table 1 provided for the exposure times established in this study. In addition, please be aware that the DAB solution is toxic and carcinogenic. Ensure that you dispose of the solution properly, in accordance with your institution’s guidelines.
13. After washing, proceed to hematoxylin counterstaining. Stain the slides with hematoxylin for 1 minute. Rinse off the excess hematoxylin with tap water 3 times. Then, wash with a destaining solution (70% ethanol and 1% HCl in water) for a few seconds to slightly reduce the intensity of the hematoxylin counterstain. Rinse with cold tap water 3times. Next, stain the slides with 0.2% ammonium hydroxide (pH 10) for 10 minutes to achieve bluing. Finish by washing twice with cold tap water and then perform a final wash with MilliQ water.
Note: Alcohol destaining and bluing steps are optional, which we preferas it shows better nucleus structure and membrane.
14. Proceed with the dehydration of FFPE slides using the following protocol:

a. 1 70% Ethanol wash for the slide for the duration of 3 minutes.
b. 1 95% Ethanol wash for the slide for the duration of 3 minutes.
c. 2 100% Ethanol washes for the slide, each wash is for the duration of 3 minutes.
d. 2 Xylene washes for the slide, each wash is for the duration of 5 minutes.
15. Mount the slides using Cytoseal. Allow the slides to dry in the fumehood overnight before starting the imaging process.

### Microscopy

Microscopy was performed using the Nikon Eclipse Ni-U Upright microscope with 40x Nikon Plan Fluor objective. Images were acquired using the Nikon DS-Fi3 camera and NIS-Elements BR software (Version 5.20.01). While image acquisition, exposure time was adjusted using the Color DS-Fi3 settings to obtain a clear definition of tissue structure. Images were acquired in the TIFF format.

**Table 1:**
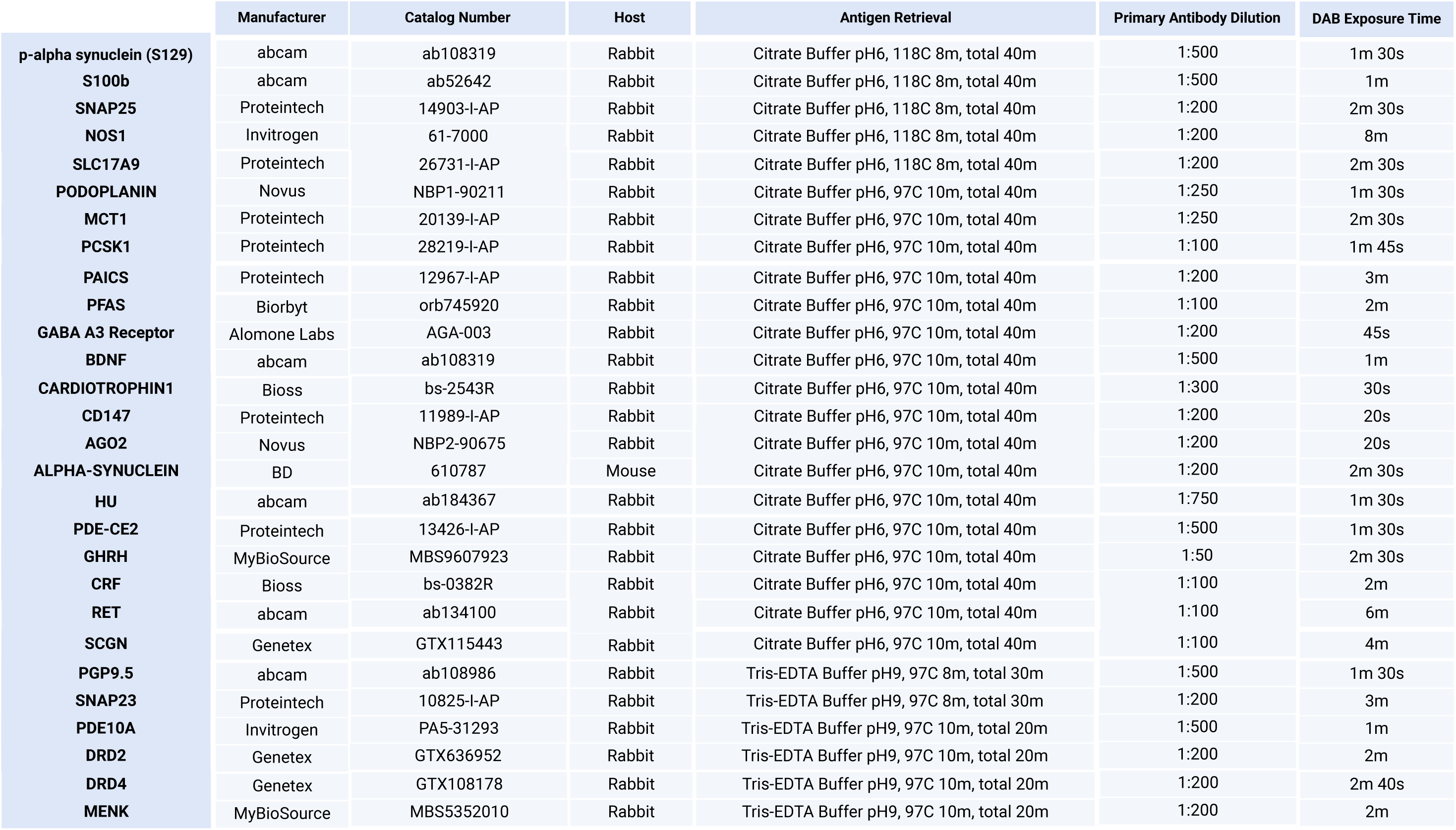
Detailed list of antibodies, dilution, antigen retrieval parameters and DAB parameters for each of the optimized antigen detection protocol.

## Results

### Detection of various ENS-associated antigens can be performed by varying antigen retrieval conditions and DAB exposure time

Using archived FFPE tissues, we performed immunohistochemical interrogation of sections of adult human small intestinal full thickness tissue. We used antibodies against 35 different antigens and were able to optimize the immunostaining against 27 antigens. The antibodies and the conditions used for detecting these antigens, which include pan-neuronal markers, glial marker, neurotransmitter encoding genes, receptors, hormones, and enzymes, in adult human small intestinal ganglia are tabulated in Table 1. We will discuss the antigens detected in defined categories and succinctly describe their relevance to ENS health and disease next.

### Neuronal markers

We first assessed a set of neuronal markers that are known to be expressed by all neurons or by a subset of neurons (**Fig 1**). Our assessment initiated with the pan-neuronal marker Hu (**Fig 1A**), which we observe labels both nuclear and cytoplasmic fractions of the cell^22–24^. After antigen retrieval, the nuclear fraction of the cell is immunostained more heavily than the cytoplasmic fraction. Next, we optimized the immunostaining of another oft-used pan-neuronal marker PGP9.5^25^, which is observed to immunolabel the entire myenteric ganglia (both nuclear and cytoplasm) (**Fig 1B**). In addition to Hu and PGP9.5, the expression of Synaptosomal-associated protein 25 (SNAP-25) was previously thought to label all neurons of the myenteric ganglia^26^. However, prior publications from our lab and others showed that this marker only labels a subset of neurons in the murine and human myenteric plexus^27,28^. Yet, since Barrenschee et al.^27^ found that the expression of this molecule, which is the target for BOTOX that is used for treatment of achalasia^29,30^, is modulated by presence of GDNF, we optimized the immunostaining parameters for this antigen. We found significant immunoreactivity of SNAP-25 in the fibers surrounding the neurons inside the ganglia and weak nuclear immunoreactivity in a subset of neurons (**Fig 1C**). While these three proteins have been studied in the gut, the expression of Phosphodiesterase 10A (PDE10A), which is a dual-substrate PDE that is highly enriched in brain^31^, has not been studied in the human ENS. We previously tested and found that PDE10A was expressed by myenteric neurons in the adult mouse small intestine^28^. With that background, we performed optimization of the immunostaining for PDE10A on human gut tissues and found that while PDE10A was not specifically expressed by neurons in the adult human gut, it was expressed by many cells within the enteric ganglia (**Fig 1D**) (and by many cells in epithelial crypts). Importantly, all the 4 neuronal markers optimized by us vary either in the nature of the antigen retrieval buffer or the conditions for antigen retrieval and DAB staining.

**Figure 1.**
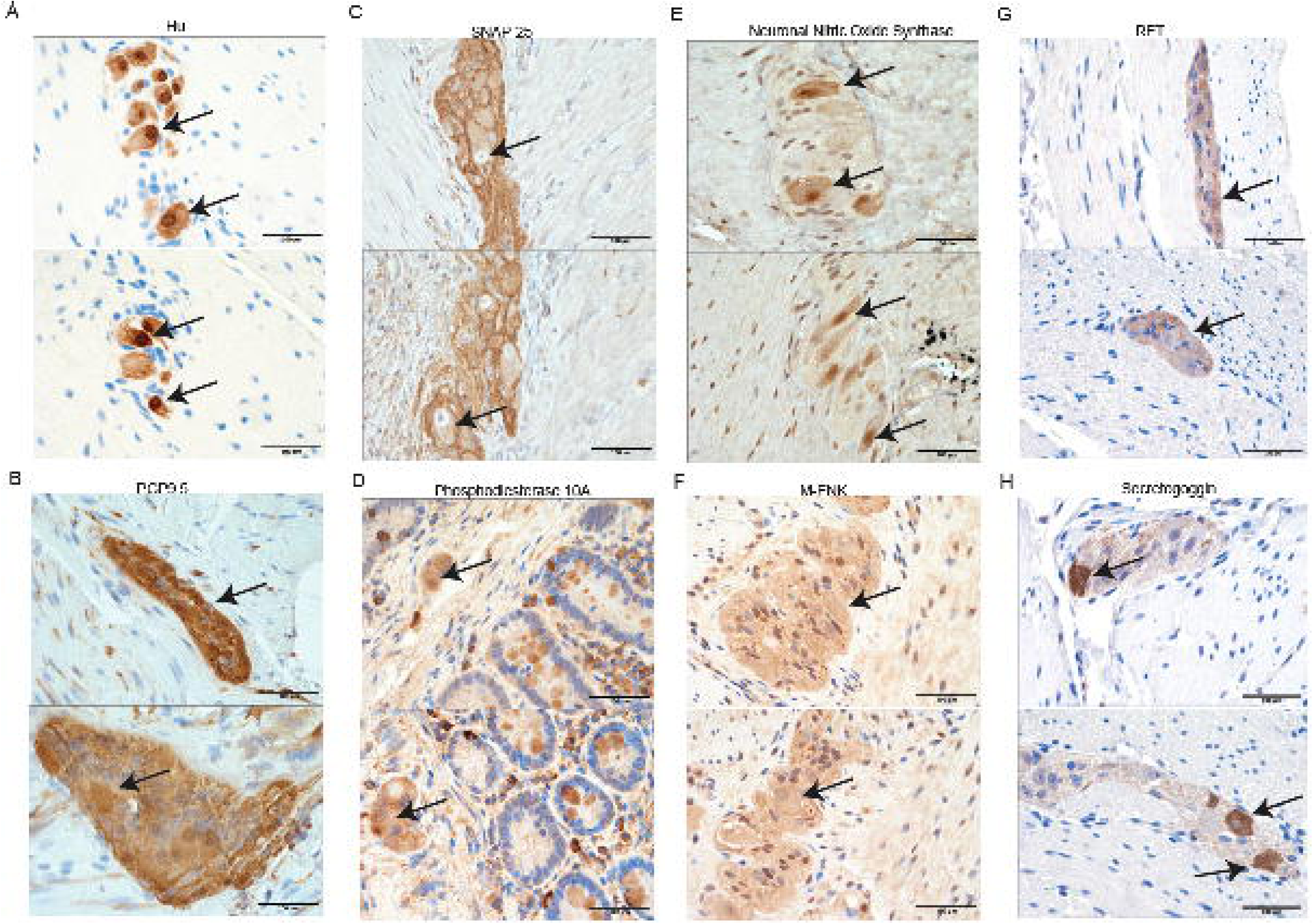
Representative images of immunohistochemical detection of important neuronal markers in the adult small intestinal myenteric ganglia. Representative images showing detection of important neuronal markers (A) Hu, (B) PGP9.5, (C) SNAP-25, (D) Phosphodiesterase 10A, (E) Neuronal Nitric Oxide Synthase (NOS1), (F) Methionine-Enkephalin (M-ENK), (G) RET, and (H) Secretogoggin that were imaged under 40X objective after optimized immunostaining on 5 μm thick formalin fixed paraffin embedded sections of adult human small intestinal tissues. Arrows in A, D, E, H mark neurons showing positive immunoreactivity. Arrows in B, F, G show cells with increased immunoreactivity in ganglia that are otherwise diffusely immunolabeled. Arrows in C show cells with nuclear immunoreactivity. Scale bar is 100 μm.

We next optimized the immunostaining of Nitric Oxide Synthase 1 (NOS1), which is expressed by inhibitory nitrergic neurons in the myenteric plexus, which is the population of neurons selectively lost in the lower esophageal sphincter in achalasia^32^. Loss of NOS1^+^ neurons has been associated with gastric and colonic dysmotility in pre-clinical models^33,34^, and thus, the optimization of the immunohistochemical staining of this marker in human small intestinal tissue is clinically significant. Our optimzed protocol shows that NOS1 is expressed only in the soma of a select subset of neurons in the myenteric ganglia (**Fig 1E**). We also observe nuclear immunostaining of NOS1, which is possibly a non-specific staining.

Enkephalins, which are endogenous opioids that signal through opioid receptors, are expressed by about a quarter of all rodent myenteric neurons^35,36^. Proenkephalin (PENK) is the precursor enkephalin which is catalytically cleaved into smaller functional enkephalins, which include the pentapeptide Methionine-enkephalin (M-ENK)^37^. M-ENK is also known as Opioid Growth Factor (OGF) which has important anti-cancer properties, and we tested whether M-ENK is expressed in the ENS. By optimizing the immunostaining for this antigen, we observed that M-ENK immunoreactivity is detected in the gut wall, and its expression is enriched in the myenteric ganglia and nuclear and cytoplasmic localization can be expressed in many myenteric cells (**Fig 1F**).

We next tested the presence of RET, the receptor for glial derived neurotrophic factor (GDNF) in the adult human ENS. RET is known to be expressed in a majority of neurons in the neonatal ENS as well as in the adult ENS^38^, and it is the target of putative therapies against irritable bowel syndrome (IBS)^39^. We recently showed that with age, the expression of RET is significantly reduced in the murine ENS and hence, here, we optimized the immunostaining for this important marker such that it may be used to assess how RET expression changes in the adult ENS of patients with myriad GI dysmotility disorders. On optimizing the expression, we found that RET expression in the adult human gut wall is restricted to the ganglia, where it exhibits sparse and light expression (**Fig 1G**). It is not readily discernable whether RET immunochemistry is restricted only to neurons within the ganglia or if it is observed in glial cells and nerve fibers.

Since we studied the limited expression of SNAP-25, we tested and optimized the detection method for Secretagoggin (SCGN), which is a calcium-sensing protein that interacts with SNAP-25 in a Ca^2+^-dependent manner to regulate membrane fusion events^40^. SCGN is a risk gene for autism spectrum disorders, and its deficiency is associated with increased susceptibility of developing colitis^40,41^. Furthermore, *Scgn* gene expression has been observed in murine and human ENS^36,42^. By optimizing the detection of SCGN in human ENS, we observe a robust immunoreactivity for this protein in the soma and nucleus of a defined subset of enteric neurons (**Fig 1H**).

### Neurotransmitter receptors

We tested a few neurotransmitter receptors and were able to optimize the protocols for the detection of three receptors, two for Dopamine and one for Gamma Amino Butyric Acid (GABA) (**Fig 2**). Dopamine receptor D2 (DRD2) expression was previously observed in the murine ileum and colon^43,44^ and DRD2 antagonists are known to have a prokinetic effect on rodent and human gut motility^45^. Our optimized protocol detects robust DRD2 expression in most cells of the myenteric ganglia which include both neurons and glial cells, but also detects cytoplasmic DRD2 expression in extra-ganglionic cells in the longitudinal and circular muscle layer (**Fig 2A**). Compared to DRD2, not much is known about expression of Dopamine Receptor D4 in the gut wall, despite being a known drug target for psychostimulants such as Clozapine that are known to affect GI motility^46,47^. Our optimized protocol detects DRD4 expression both in a subset of myenteric neurons, as well as in the surrounding muscle cells of the circular and longitudinal muscle layers (**Fig 2B**). DRD4 expression appears to be cytoplasmic only in most of the cells stained by this antibody, but in a subset of cells, strong immunoreactivity is observed in both cytoplasm and nucleus. In addition to Dopaminergic receptors, we tested the protocols for optimization of various receptors for GABA and were able to optimize the protocols to allow the robust detection of GABA Receptor A3 (GABRA3), which is a target for benzodiazepines. GABRA3 was previously shown to be expressed in murine colonic ENS^48^, and here, by optimizing the protocol for its detection in adult human ENS, we observe robust expression of GABRA3 in a subset of myenteric neurons while also detecting lower level expression of this receptor in non-neuronal cells within and outside the myenteric ganglia (**Fig 2C**). While it is unclear why expression of GABRA3 is observed in nucleus of many cells, the database at human protein atlas shows nucleoplasm localization of this protein.

**Figure 2.**
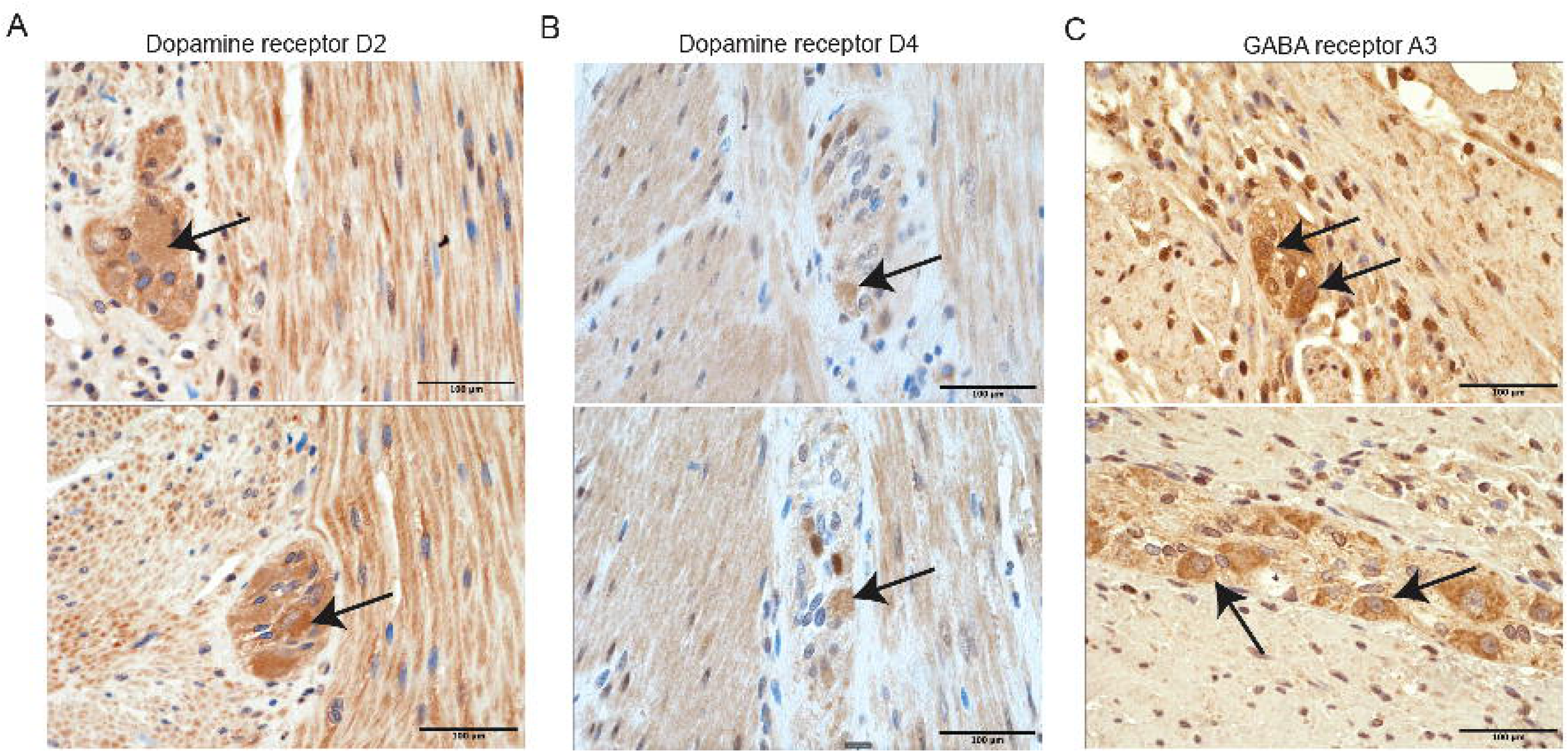
Representative images of immunohistochemical detection of important neurotransmitter receptors in the adult small intestinal myenteric ganglia. Representative images showing detection of important neuronal markers (A) Dopamine Receptor D2, (B) Dopamine Receptor D4, and (C) GABA receptor A3 that were imaged under 40X objective after optimized immunostaining on 5 μm thick formalin fixed paraffin embedded sections of adult human small intestinal tissues. Arrows show cells with increased immunoreactivity in ganglia that are otherwise contain low diffuse immunoreactivity. Scale bar is 100 μm.

### Hormones

We tested whether we could optimize the expression of neuronal hormones in the ENS and were able to optimize the detection of two important neuronal hormones (**Fig 3**). Corticotrophin releasing hormone (CRH) is known to be expressed by enteric neurons in addition to its known expression by hypothalamic neurons^49,50^. CRH release is known to be associated with various stress-associated dysmotility disorders, which makes it an important marker to establish for the adult human ENS. Here, we optimize the immunochemical detection protocol for CRH to observe significant and robust nuclear and cytoplasmic expression of this hormone by a subset of adult human small intestinal myenteric neurons (**Fig 3A**). Similar to CRH, Growth Hormone Releasing Hormone (GHRH, Somatoliberin) is expressed by hypothalamic neurons^51^. We recently detected the expression of GHRH in a subset of adult murine myenteric neurons that are derived from the mesoderm (and annotated as mesoderm-derived enteric neurons, MENs^28^). Here, we optimized the detection protocols and observed that GHRH is robustly localized in the nucleus and cytoplasm of a subset of adult human small intestinal myenteric neurons (**Fig 3B**), and as with CRH, the possible release of this hormone into the myenteric ganglia and in the gut wall produces a diffuse and low level immunoreactivity for this hormone in GHRH-non-expressing neurons. Both the hormones require similar antigen retrieval and DAB exposure settings.

**Figure 3.**
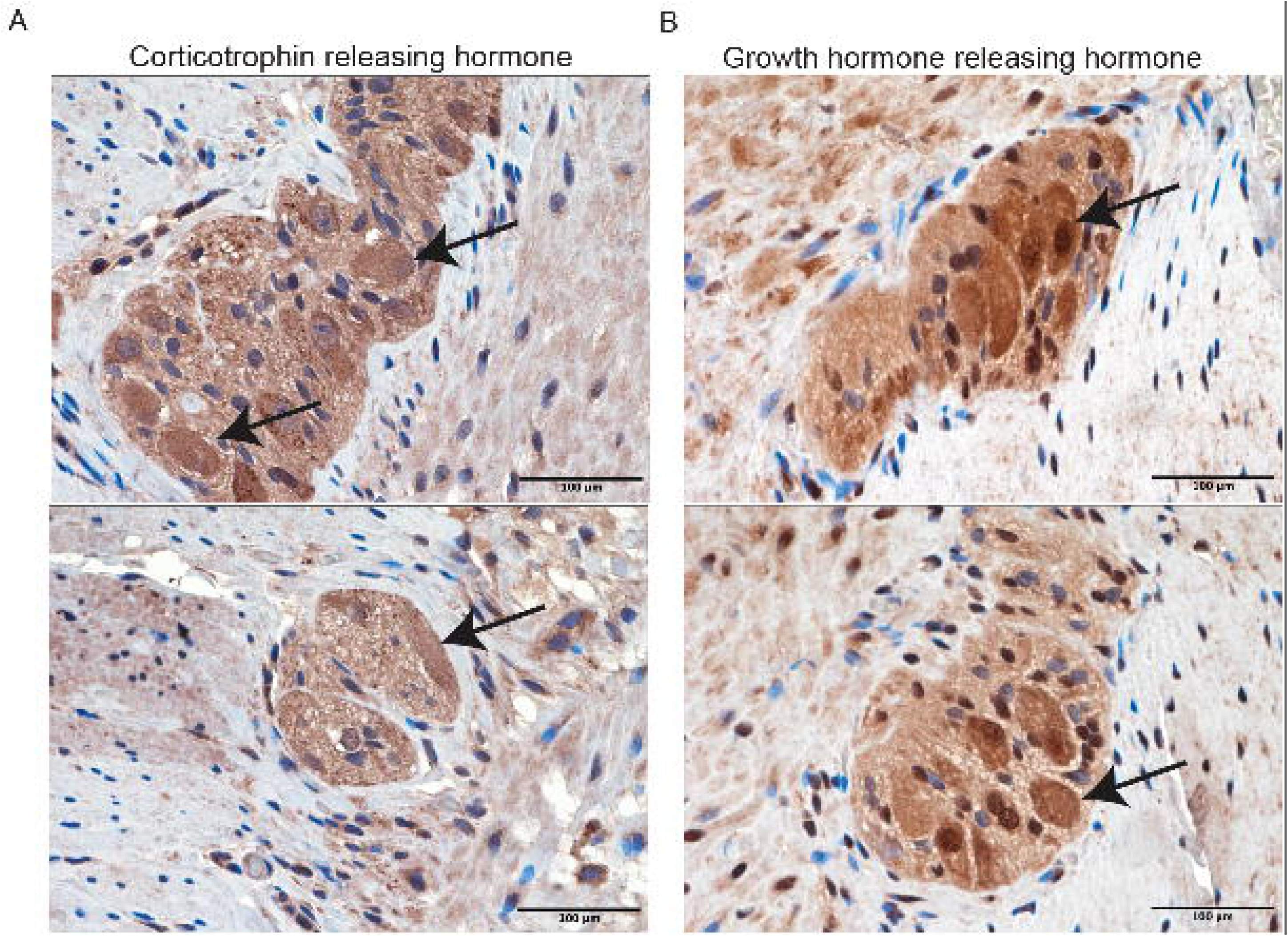
Representative images of immunohistochemical detection of important hormones in the adult small intestinal myenteric ganglia. Representative images showing detection of important neuronal markers (A) Corticotrophin Releasing Hormone, and (B) Growth Hormone Releasing Hormone that were imaged under 40X objective after optimized immunostaining on 5 μm thick formalin fixed paraffin embedded sections of adult human small intestinal tissues. Arrows show cells with increased immunoreactivity in ganglia that are otherwise contain low diffuse immunoreactivity. Scale bar is 100 μm.

### Alpha-synuclein

Recent gathering evidence suggests that the pathological misfolded species of the protein alpha synuclein that drives the etiology of Parkinson’s’ disease (PD) originates in the gut, from where it is trafficked to the central nervous system to cause the loss of dopaminergic neurons in the substantia nigra thus manifesting the behavioral and motor symptoms of the disease^52–56^. Prior reports have suggested that the normal and pathological alpha-synuclein can be distinguished by the presence of phosphorylated Serine at residue 129 (pS129), as this phosphorylation was observed to occur in the pathological lewy bodies associated with the disease^57^. However, the relevance of the phosphorylated form in the ENS to PD is unclear^58^. Thus, it is important to optimize the detection protocols for both the total and pS129 forms of alpha-synuclein in the adult human gut, so that investigators can use this methodology for accurate assessment of the presence of these proteins in the gut wall of patients with and without PD. Here, with optimization which involved two distinct antigen retrieval conditions for the two proteins, we show evidence that both the normal (**Fig 4A**) and the pS129 forms (**Fig 4B**) of alpha-synuclein can be detected in the myenteric neurons of patients without PD. Both forms of the protein can be detected in the nucleus and cytoplasm of immunostained cells.

**Figure 4.**
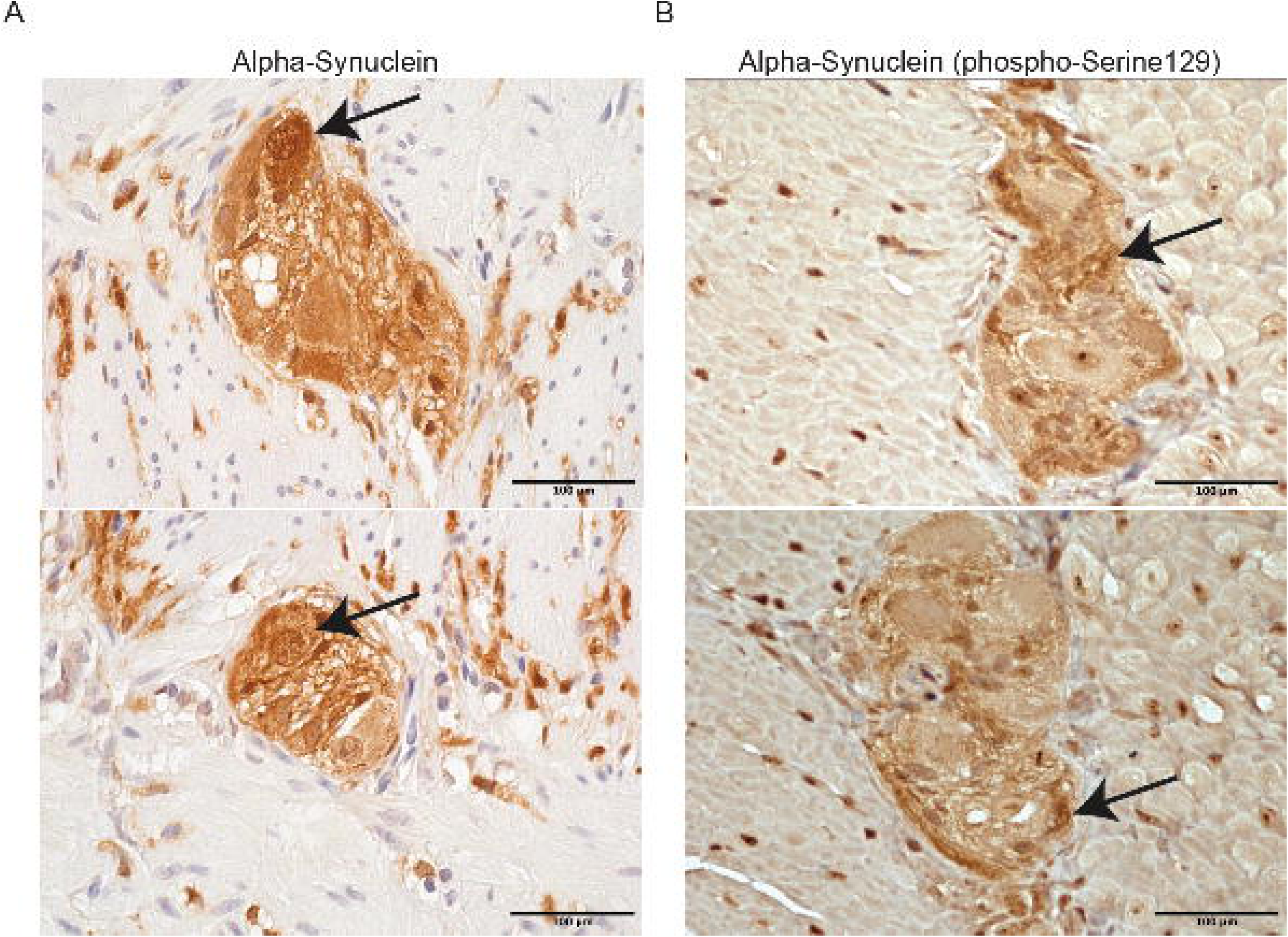
Representative images of immunohistochemical detection of alpha-synuclein in the adult small intestinal myenteric ganglia. Representative images showing detection of (A) total alpha-synuclein, and (B) phosphorylated at Serine 129 form of alpha-synuclein (pS129-a-syn) that were imaged under 40X objective after optimized immunostaining on 5 μm thick formalin fixed paraffin embedded sections of adult human small intestinal tissues. Arrows in A show cells with increased immunoreactivity in ganglia that are otherwise contain low diffuse immunoreactivity, and arrows in B show regions within the ganglia that show increased abundance of pS129-a-syn. Scale bar is 100 μm.

### Purine anabolism

Purine anabolism occurs through the de novo pathway or the salvage pathway^59^, and in the small intestine, the de novo pathway of purine anabolism – which is catalyzed by 6 enzymes^60^ – is favored^59^. Disorders of purine metabolism result in several disease conditions, which include irritable bowel syndrome^61^ and autism spectrum disorders that are associated with GI dysmotility^62,63^. Thus, given the importance of purine metabolism to GI conditions, we optimized the detection of 2 important enzymes of the de novo purine anabolic pathway – PAICS (Phosphoribosylaminoimidazolesuccinocarboxamide Synthase) and PFAS (Phosphoribosylformylglycinamidine Synthase) in the adult human small intestinal ENS and found that both enzymes are robustly expressed in a subset of human myenteric neurons (**Fig 5**). PAICS (**Fig 5A**) shows a predominantly cytoplasmic expression which is restricted to the myenteric ganglia, while PFAS shows predominantly nuclear localization (**Fig 5B**) which is also observed in mucosal cells of the small intestine.

**Figure 5.**
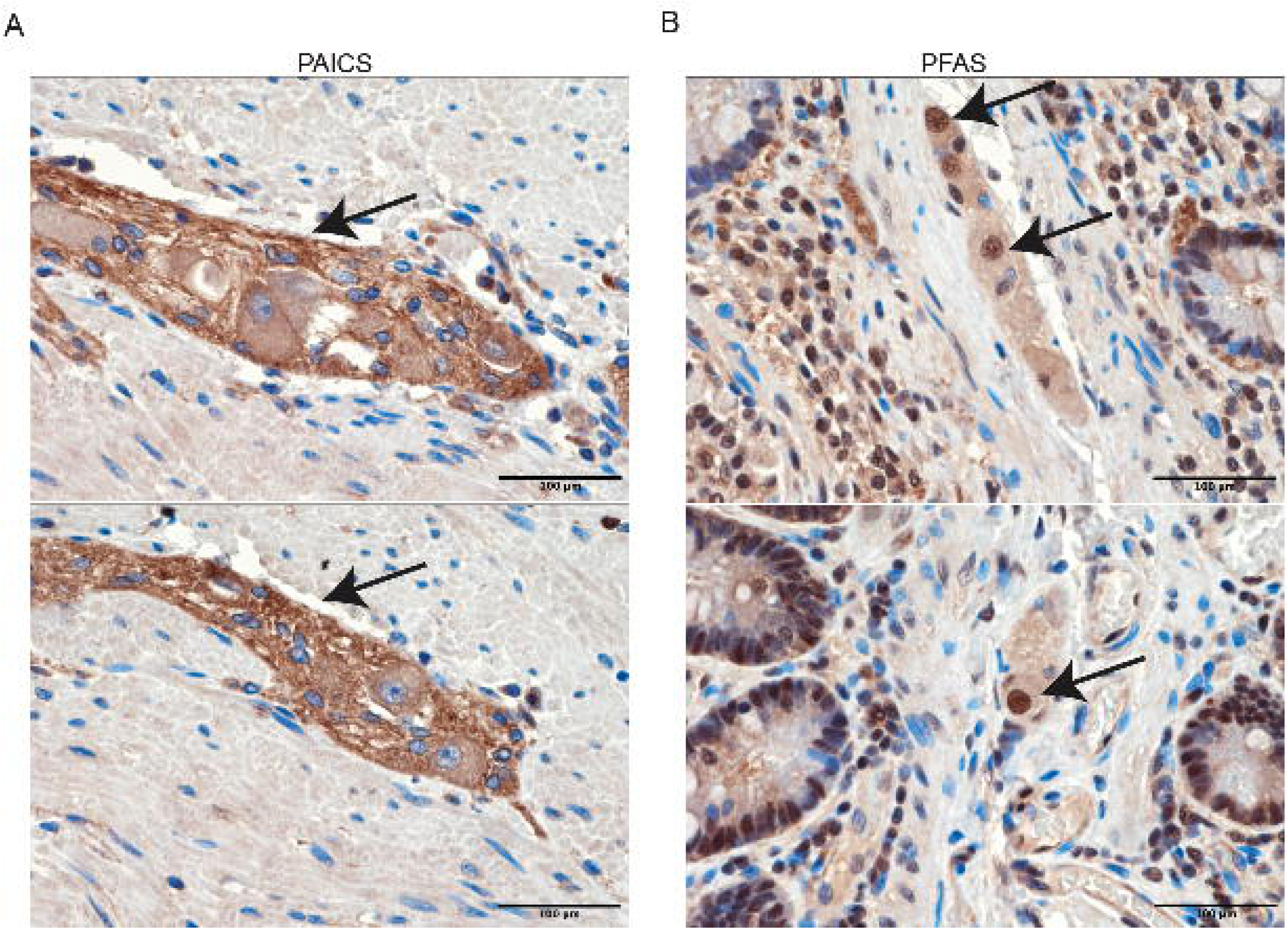
Representative images of immunohistochemical detection of enzymes involved in de novo purine biosynthesis in the adult small intestinal myenteric ganglia. Representative images showing detection of enzymes (A) PAICS, and (B) PFAS that were imaged under 40X objective after optimized immunostaining on 5 μm thick formalin fixed paraffin embedded sections of adult human small intestinal tissues. Arrows in (A) show regions in the ganglia with increased immunoreactivity amongst other regions that otherwise contain low diffuse immunoreactivity, and arrows in B show ganglionic cells with nuclear localization of PFAS. Scale bar is 100 μm.

### Cellular metabolism

Metabolic and mitochondrial disorders are associated with increased prevalence of GI dysmotility disorders^64,65^. Preclinical models have tested and found the causative nature of metabolic and mitochondrial imbalances on ENS structure and gut function^66,67^. Given this, it is important to optimize where important markers involved in cellular metabolism are located in the adult human ENS. First, we optimized the expression of a monocarboxylate transporter 1 (MCT; SLC16A1) which helps import monocarboxylate acids such as Lactate, Pyruvate which are known substrates for oxidative phosphorylation into the cell^68^. In skeletal muscles, MCT1 plays a significant role in the Lactate Shuttle into the cell to promote mitochondrial biogenesis^68^. Our optimized protocol for detecting MCT1 in adult ENS shows that MCT1 is expressed by a subset of enteric neurons (**Fig 6A**). The localization of this protein appears to be membrane bound, as would be expected from a transporter protein. Next, we assessed whether CD147 or Basigin – which regulates the expression of MCT1 and Lactate Shuttle^69^ – is also expressed in adult human ENS. MCT1 and other monocarboxylate transporters also bind to CD147, as CD147 interaction with MCT1 stabilizes both binding partners as they are deployed to the cell surface^70^. Importantly, both MCT and CD147 proteins are targets in putative cancer therapies that target lactate metabolism^70^. Our optimized protocol shows robust and enriched expression and putative membrane-associated localization of CD147 in adult human myenteric neurons (**Fig 6B**) – similar to our observations with MCT1. In addition, some cells around the myenteric ganglia also show nuclear localization. and the Since Lactate importer MCT1, which is crucial for promoting mitochondrial function, is observed to be expressed by only a subset of neurons, we next tested whether a pan-mitochondrial protein of clinical significance is also expressed by a subset of myenteric neurons. By optimizing a detection protocol with antibodies against the mitochondrial M2 antigen (PDC-E2; Pyruvate Dehydrogenase Complex – E2 subunit; gene name *Dlat*) – which is the main antigen targeted in patients with Primary Biliary Cirrhosis^71^, we found that a subset of enteric neurons are highly enriched in M2 antigen and hence mitochondria (**Fig 6C**). These mitochondria-laden myenteric neurons are the sole cells that show significant abundance of mitochondria in the gut wall, and our results based on our optimized detection protocol help provide a rationale for why metabolic and mitochondrial diseases overwhelmingly cause a loss of healthy GI motility in patients.

**Figure 6.**
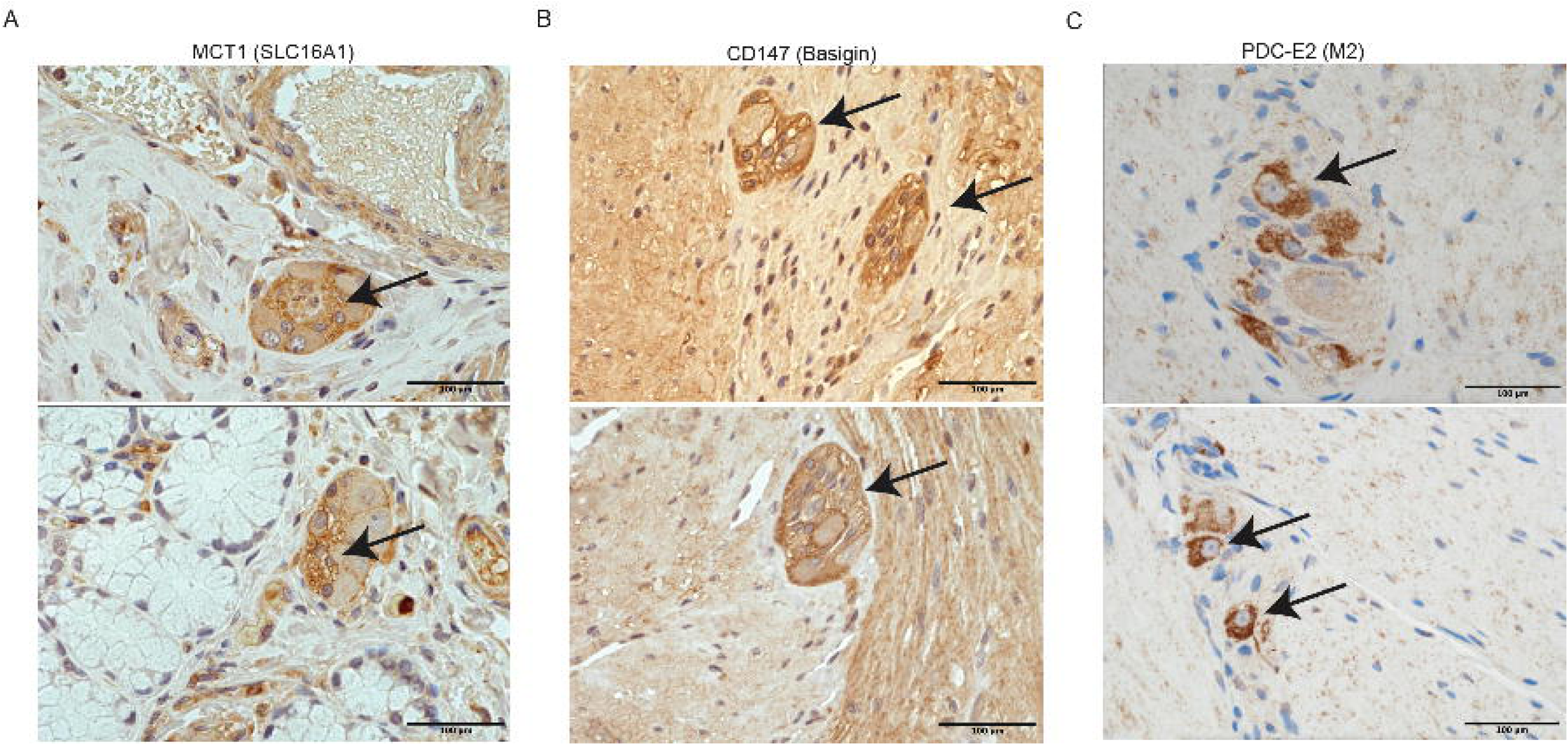
Representative images of immunohistochemical detection of metabolically important proteins in the adult small intestinal myenteric ganglia. Representative images showing detection of (A) Lactate transporter MCT1 (SLC16A1), (B) CD147 or Basigin, and (C) mitochondrial antigen M2 also known as PDC-E2, that were imaged under 40X objective after optimized immunostaining on 5 μm thick formalin fixed paraffin embedded sections of adult human small intestinal tissues. Arrows in A and B show putative membrane-bound immunoreactivity in myenteric cells, and arrows in C show that a subset of myenteric neurons alone are enriched in the M2 antigen. Scale bar is 100 μm.

### Other markers

In addition to these markers, we also optimized the detection protocols for the following clinically or biologically significant markers. Deficiency in Proprotein convertase subtilisin/kexin type 1 (PCSK1) is a congenital anomaly which is associated with significant GI dysmotility^72^. We optimized the detection of PCSK1 in the adult human ENS and observed that PCSK1 immunoreactivity in the gut wall is robustly present in the nucleus and cytoplasm of cells (putative neurons) of the myenteric ganglia, with surrounding cells exhibiting low level cytoplasmic but may exhibit strong nuclear immunoreactivity (**Fig 7A**).

**Figure 7.**
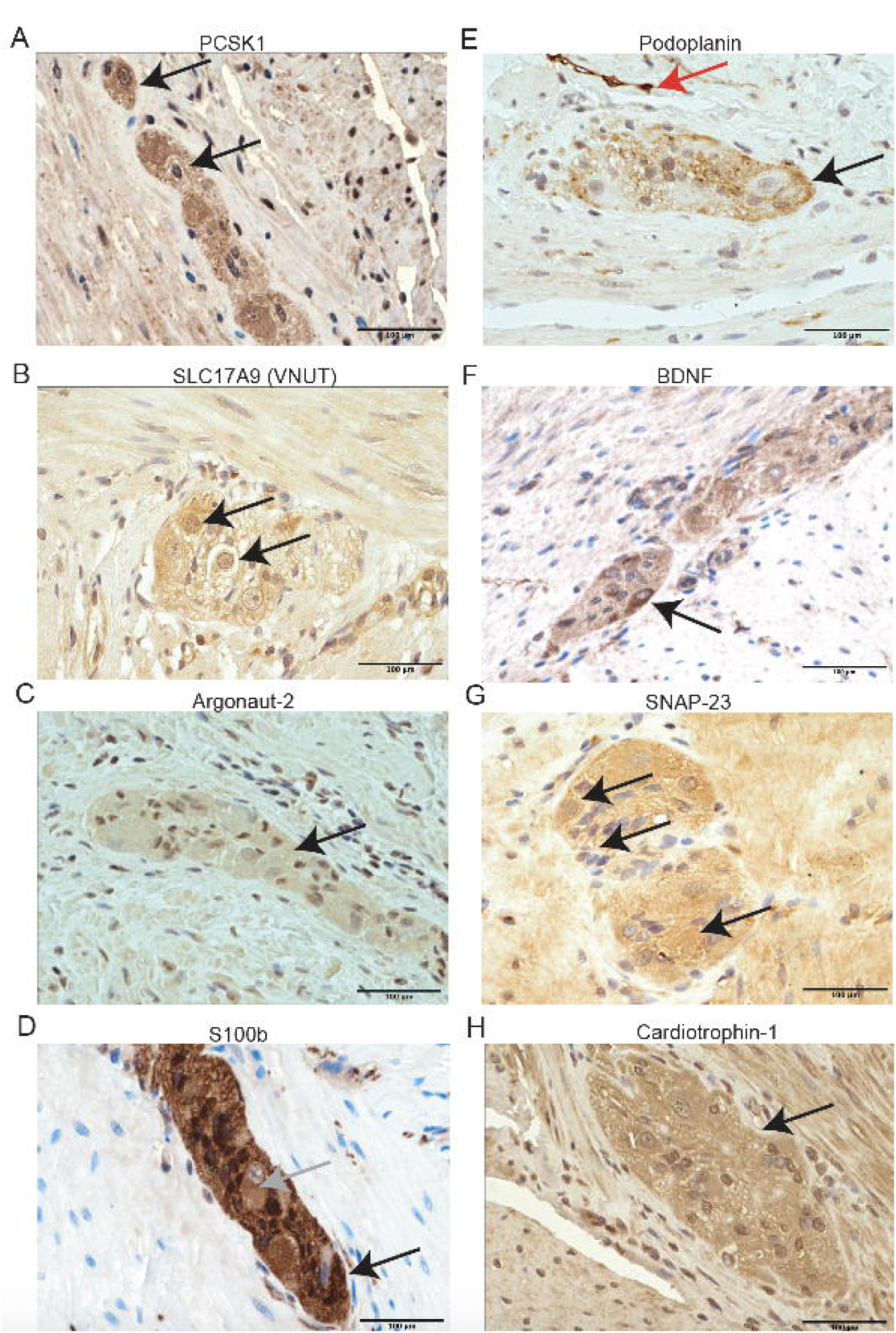
Representative images of immunohistochemical detection of clinically and pathologically important proteins in the adult small intestinal myenteric ganglia. Representative images showing detection of (A) PCSK1, (B) SLC17A9 (VNUT), (C) Argonaut-2, (D) S100b, (E) Podoplanin, (F) BDNF, (G) SNAP-23, and (H) Cardiotrophin-1 that were imaged under 40X objective after optimized immunostaining on 5 μm thick formalin fixed paraffin embedded sections of adult human small intestinal tissues. Arrows in A and F show neurons that show marked increase in abundance of the markers compared to relatively lower levels in other ganglionic cells. Arrows in B show cells with nuclear labeling and increased abundance of marker relative to other cells in the ganglion. Arrows in C, G, H show diffuse ganglionic immunoreactivity. Black arrow in D shows strongly positive glial cell and gray arrow in this panel shows relatively lower immunoreactivity in neurons. Black arrow in E shows putative membrane immunoreactivity in myenteric neurons and red arrow in the same panel shows strong immunoreactivity in lymphatic endothelium. Scale bar is 100 μm.

SLC17A9 (Vesicular Nucleotide Transporter; VNUT) is a that plays a central role in vesicular storage of nucleotide in purine-secreting cells. Since we earlier noted purine anabolism in myenteric neurons, and we have previously detected SLC17A9 gene and protein expression in murine ENS, we optimized the detection of this protein in adult human ENS^28^. We observed nuclear and cytoplasmic localization of SLC17A9 in the myenteric neurons, but also its expression in the surrounding gut musculature (**Fig 7B**).

Recent studies identify a significant role for extracellular vesicles (EVs) in the regulation of biology of health and disease^73^. The protein Argonaut-2 (AGO2) is a key component of the RNA-induced silencing complex (RISC) present within EVs that mediates downregulation of mRNA by microRNAs^73^. Thus, given its potential importance in health and disease, we optimized the detection of AGO2 in the adult human gut wall and found that both myenteric ganglia and surrounding cells showed moderate cytoplasmic and strong nuclear localization of this protein (**Fig 7C**). AGO2 abundance was slightly higher in myenteric ganglia which may suggest that the ganglia are the source of the AGO2-EVs, which are released into the surrounding tissues.

The protein S100B has been often used as a ganglionic marker in pathological interrogation of gut wall, while it is used primarily as a glial marker in the preclinical models^74–76^. We optimized the detection parameters for this antibody and found that nuclear and cytoplasmic immunoreactivity of S100B was very high in the myenteric ganglia with both neurons and glial cells expressing this protein (**Fig 7D**). However, glial cells expressed this marker much more robustly than neurons whose cytoplasmic immunoreactivity was a magnitude lower than in glial cells.

Podoplanin is an approximate 38-kDa membrane protein with several conserved O-glycosylation sites. Heavily O-glycosylated mucoproteins are counter receptors for selectins that mediate adhesion of inflammatory cells, and as a result Podoplanin expression in intestinal lymphatics has been observed to be important in the pathogenesis of inflammatory bowel diseases^77^. Our recent study identified that Podoplanin is expressed by mesoderm-derived enteric neurons, and thus we tested whether we can find the expression of this important marker in adult myenteric neurons. Our optimized detection allows us to detect putative membrane-bound immunoreactivity of this protein in the myenteric neurons, along with detecting strong immunoreactivity for this protein in putative lymphatic vasculature (**Fig 7E**).

Brain-derived neurotrophic factor (BDNF) is an important neurotrophin that is expressed in the ENS and has been shown to be implicated in GI dysmotility, IBS^78–81^. Given its importance to GI health and disease, we optimized the detection protocol and found that BDNF protein expression is highly enriched in the cytoplasm of a subset of myenteric neurons (**Fig 7F**). Since BDNF is a secreted protein, we reason its detection in other myenteric cells is a result of the detection of the secreted form.

In addition to SNAP-25, another synaptosomal associated protein 23 (SNAP-23) is also expressed by neurons and other cells that perform exocrine and endocrine functions^82–84^. Overexpression of alpha-synuclein, which occurs in Parkinson’s’ Disease, is known to reduce the expression of SNAP-23^85^. Since we find significant expression of alpha-synuclein in the myenteric plexus, and overexpression of alpha-synuclein in a preclinical setting is known to cause GI dysmotility^86^, we optimized the detection of SNAP-23 in the adult ENS, and found that this protein is the cytoplasm of both myenteric neurons as well as in cells of surrounding gut musculature (**Fig 7G**). The optimization of this protocol along with that for alpha-synuclein provides us with important tools through which the cellular and molecular aberrations in the ENS of PD patients can be assessed.

Cardiotrophin-1 is a member of the IL-6 family of cytokines, and it is responsible for inducing cardiac hypertrophy^87^. Apart from its role in regulating cardiac health, Cardiotrophin-1 also regulates intestinal immunity as it prevents the development of intestinal inflammation in a chemically-induced model of colitis^88^. Given the importance of neuro-immune crosstalk in regulation of intestinal immunity^89^, we optimized the detection of this important secreted cytokine in the human gut wall and found that Cardiotrophin-1 is abundantly detected in the nucleus and cytoplasm of cells of myenteric ganglia of adult human small intestine (**Fig 7H**). The surrounding cells of the gut musculature also show detection of this cytokine, which we assume are the secreted forms released by the myenteric ganglion.

## Discussion

Understanding the pathobiology of GI dysmotility disorders requires us to study how the enteric nervous system (ENS) and surrounding cells are impacted at a structural and molecular level. Without this fundamental pathobiological understanding of GI dysmotility, our attempts at simulating disease in preclinical models contain a significant gap – as how the tissue level changes in preclinical models correlate with actual changes in patients remain unknown. This study that provides optimized methods for interrogation of human ENS tissue using molecular markers that are clinically relevant provides an important bridge through which the relevance of preclinical studies to clinical histopathology can be assessed.

Recent studies on performing assessment of how ENS is impacted in clinical tissues have focused on using freshly isolated full thickness whole mount tissues of longitudinal muscle – myenteric plexus (LM-MP) preparations from fetal and post-natal human gut^90,91^. These studies provide a 3-dimensional (3D) view of the ENS, which provide for better enumeration and interrogation of ENS structure. However, since these studies require freshly harvested tissues, they can be performed only on prospectively captured tissues. These approaches are thus not applicable to the thousands and potentially millions of retrospectively captured human gut specimens in FFPE tissue blocks^19^ from patients without and with motility disorders. Further, the advent of novel computational tools such as CODA allow for the 3D reconstruction of sectioned FFPE tissue blocks, thus allowing for accurate representation and interrogation of sectioned and immunostained tissues existing in FFPE tissue blocks^92^. The use of such tools on retrospectively captured FFPE tissue blocks provides the ideal way forward through which significant immunohistochemical assessment of ENS tissue from patients can be performed to learn and diagnose how this tissue is impacted in patients with GI dysmotility conditions. The optimization of methods and results that we report in this study are thus the first and important step through which comprehensive assessment of ENS and other gut cells can be performed.

This report, which describes the initial set of methods optimized to detect clinically important antigens in the adult human ENS, is the foundation for an openly accessible online platform where methods and image data which will be made freely available to the GI and Pathology community. Such a resource will be valuable for catalyzing the translational science and the diagnostic science of understanding GI dysmotility disorders in patients.

## Acknowledgements

We acknowledge Dr. Philippa Seika, Dr. Trisha Pasricha, and Mrs. Vilmonse (Eva) Csizmadia for their help. This work was supported by funding from NIA R01AG66768 (SK), and the Diacomp Foundation (Pilot award Augusta University).

## References

1 Sperber, A. D. et al. Worldwide Prevalence and Burden of Functional Gastrointestinal Disorders, Results of Rome Foundation Global Study. Gastroenterology 160, 99–114.e113 (2021). 10.1053/j.gastro.2020.04.014

2 Leigh, S.-J. et al. The impact of acute and chronic stress on gastrointestinal physiology and function: a microbiota–gut–brain axis perspective. The Journal of Physiology 601, 4491–4538 (2023). 10.1113/JP281951

3 Sharkey, K. A. & Mawe, G. M. The enteric nervous system. Physiological Reviews 103, 1487–1564 (2023). 10.1152/physrev.00018.2022

4 Obata, Y. & Pachnis, V. The Effect of Microbiota and the Immune System on the Development and Organization of the Enteric Nervous System. Gastroenterology 151, 836–844 (2016). 10.1053/j.gastro.2016.07.044

5 Nezami, B. G. & Srinivasan, S. Enteric nervous system in the small intestine: pathophysiology and clinical implications. Curr Gastroenterol Rep 12, 358–365 (2010). 10.1007/s11894-010-0129-9

6 Mourad, F. H. & Saade, N. E. Neural regulation of intestinal nutrient absorption. Prog Neurobiol 95, 149–162 (2011). 10.1016/j.pneurobio.2011.07.010

7 Mazzuoli-Weber, G. & Schemann, M. Mechanosensitivity in the enteric nervous system. Front Cell Neurosci 9, 408 (2015). 10.3389/fncel.2015.00408

8 Gariepy, C. E. Intestinal motility disorders and development of the enteric nervous system. Pediatr Res 49, 605–613 (2001). 10.1203/00006450-200105000-00001

9 Furness, J. B. & Sanger, G. J. Intrinsic nerve circuits of the gastrointestinal tract: identification of drug targets. Curr Opin Pharmacol 2, 612–622 (2002). 10.1016/s1471-4892(02)00219-9

10 Furness, J. B. The enteric nervous system: normal functions and enteric neuropathies. Neurogastroenterol Motil 20 **Suppl 1**, 32–38 (2008). 10.1111/j.1365-2982.2008.01094.x

11 Bayliss, W. M. & Starling, E. H. The movements and innervation of the small intestine. J Physiol 24, 99–143 (1899). 10.1113/jphysiol.1899.sp000752

12 Heuckeroth, R. O. Hirschsprung disease -integrating basic science and clinical medicine to improve outcomes. Nature reviews. Gastroenterology & hepatology 15, 152–167 (2018). 10.1038/nrgastro.2017.149

13 Badner, J. A., Sieber, W. K., Garver, K. L. & Chakravarti, A. A genetic study of Hirschsprung disease. Am J Hum Genet 46, 568–580 (1990).

14 Francis, D. L. & Katzka, D. A. Achalasia: update on the disease and its treatment. Gastroenterology 139, 369–374 (2010). 10.1053/j.gastro.2010.06.024

15 Boeckxstaens, G. E., Zaninotto, G. & Richter, J. E. Achalasia. Lancet 383, 83–93 (2014). 10.1016/S0140-6736(13)60651-0

16 McMahan, Z. H. et al. Systemic sclerosis gastrointestinal dysmotility: risk factors, pathophysiology, diagnosis and management. Nat Rev Rheumatol (2023). 10.1038/s41584-022-00900-6

17 Quigley, E. M. M. Constipation in Parkinson’s Disease. Seminars in Neurology 43, 562–571 (2023). 10.1055/s-0043-1771457

18 Grover, M. et al. Cellular changes in diabetic and idiopathic gastroparesis. Gastroenterology 140, 1575–1585 e1578 (2011). 10.1053/j.gastro.2011.01.046

19 Eiseman, E. & Haga, S. Handbook of human tissue sources: a national resource of human tissue samples. (Rand Santa Monica, 1999).

20 Baker, M. Biorepositories: Building better biobanks. Nature 486, 141–146 (2012). 10.1038/486141a

21 Creech, M. K., Wang, J., Nan, X. & Gibbs, S. L. Superresolution Imaging of Clinical Formalin Fixed Paraffin Embedded Breast Cancer with Single Molecule Localization Microscopy. Sci Rep 7, 40766 (2017). 10.1038/srep40766

22 Liu, M. T., Kuan, Y. H., Wang, J., Hen, R. & Gershon, M. D. 5-HT4 receptor-mediated neuroprotection and neurogenesis in the enteric nervous system of adult mice. J Neurosci 29, 9683–9699 (2009). 10.1523/JNEUROSCI.1145-09.2009

23 Desmet, A.-S., Cirillo, C. & Vanden Berghe, P. Distinct subcellular localization of the neuronal marker HuC/D reveals hypoxia-induced damage in enteric neurons. Neurogastroenterology & Motility 26, 1131–1143 (2014). 10.1111/nmo.12371

24 Bernardini, N. et al. Immunohistochemical analysis of myenteric ganglia and interstitial cells of Cajal in ulcerative colitis. J Cell Mol Med 16, 318–327 (2012). 10.1111/j.1582-4934.2011.01298.x

25 Eisenman, S. T. et al. Distribution of TMEM100 in the mouse and human gastrointestinal tract--a novel marker of enteric nerves. Neuroscience 240, 117–128 (2013). 10.1016/j.neuroscience.2013.02.034

26 Muller, P. A. et al. Microbiota-modulated CART(+) enteric neurons autonomously regulate blood glucose. Science 370, 314–321 (2020). 10.1126/science.abd6176

27 Barrenschee, M. et al. SNAP-25 is abundantly expressed in enteric neuronal networks and upregulated by the neurotrophic factor GDNF. Histochemistry and Cell Biology 143, 611–623 (2015). 10.1007/s00418-015-1310-x

28 Kulkarni, S. et al. Age-associated changes in lineage composition of the enteric nervous system regulate gut health and disease. eLife 12, RP88051 (2023). 10.7554/eLife.88051

29 Schiavo, G. et al. Botulinum neurotoxins serotypes A and E cleave SNAP-25 at distinct COOH-terminal peptide bonds. FEBS Letters 335, 99–103 (1993). 10.1016/0014-5793(93)80448-4

30 Pasricha, P. J. et al. Intrasphincteric botulinum toxin for the treatment of achalasia. N Engl J Med 332, 774–778 (1995). 10.1056/nejm199503233321203

31 Xie, Z. et al. Cellular and subcellular localization of PDE10A, a striatum-enriched phosphodiesterase. Neuroscience 139, 597–607 (2006). 10.1016/j.neuroscience.2005.12.042

32 Mearin, F., et al. Patients with achalasia lack nitric oxide synthase in the gastro-oesophageal junction. European Journal of Clinical Investigation 23, 724–728 (1993). 10.1111/j.1365-2362.1993.tb01292.x

33 Chandrasekharan, B. et al. Colonic motor dysfunction in human diabetes is associated with enteric neuronal loss and increased oxidative stress. Neurogastroenterology & Motility 23, 131–e126 (2011). 10.1111/j.1365-2982.2010.01611.x

34 Gangula, P. R. et al. Impairment of nitrergic system and delayed gastric emptying in low density lipoprotein receptor deficient female mice. Neurogastroenterol Motil 23, 773–e335 (2011). 10.1111/j.1365-2982.2011.01695.x

35 Furness, J. B., Costa, M. & Miller, R. J. Distribution and projections of nerves with enkephalin-like immunoreactivity in the guinea-pig small intestine. Neuroscience 8, 653–664 (1983). 10.1016/0306-4522(83)90001-5

36 Hamnett, R. et al. Regional cytoarchitecture of the adult and developing mouse enteric nervous system. Current Biology 32, 4483–4492.e4485 (2022). 10.1016/j.cub.2022.08.030

37 Das, K. P., Hong, J.-S. & Sanders, V. M. Ultralow concentrations of proenkephalin and [met5]-enkephalin differentially affect IgM and IgG production by B cells. Journal of Neuroimmunology 73, 37–46 (1997). 10.1016/S0165-5728(96)00165-8

38 Shepherd, A. et al. RET Signaling Persists in the Adult Intestine and Stimulates Motility by Limiting PYY Release From Enteroendocrine Cells. Gastroenterology 166, 437–449 (2024). 10.1053/j.gastro.2023.11.020

39 Russell, J. P. et al. Exploring the Potential of RET Kinase Inhibition for Irritable Bowel Syndrome: A Preclinical Investigation in Rodent Models of Colonic Hypersensitivity. J Pharmacol Exp Ther 368, 299–307 (2019). 10.1124/jpet.118.252973

40 Liu, Z. et al. SCGN deficiency is a risk factor for autism spectrum disorder. Signal Transduction and Targeted Therapy 8, 3 (2023). 10.1038/s41392-022-01225-2

41 Sifuentes-Dominguez, L. F. et al. SCGN deficiency results in colitis susceptibility. eLife 8, e49910 (2019). 10.7554/eLife.49910

42 Drokhlyansky, E. et al. The Human and Mouse Enteric Nervous System at Single-Cell Resolution. Cell 182, 1606–1622.e1623 (2020). 10.1016/j.cell.2020.08.003

43 Li, Z. S., Schmauss, C., Cuenca, A., Ratcliffe, E. & Gershon, M. D. Physiological Modulation of Intestinal Motility by Enteric Dopaminergic Neurons and the D 2 Receptor: Analysis of Dopamine Receptor Expression, Location, Development, and Function in Wild-Type and Knock-Out Mice. The Journal of Neuroscience 26, 2798–2807 (2006). 10.1523/jneurosci.4720-05.2006

44 Auteri, M., Zizzo, M. G., Amato, A. & Serio, R. Dopamine induces inhibitory effects on the circular muscle contractility of mouse distal colon via D1-and D2-like receptors. Journal of Physiology and Biochemistry 73, 395–404 (2016). 10.1007/s13105-017-0566-0

45 Tonini, M. et al. Clinical implications of enteric and central D2 receptor blockade by antidopaminergic gastrointestinal prokinetics. Alimentary Pharmacology & Therapeutics 19, 379–390 (2004). 10.1111/j.1365-2036.2004.01867.x

46 Di Ciano, P., Grandy, D. K. & Le Foll, B. Dopamine D4 receptors in psychostimulant addiction. Adv Pharmacol 69, 301–321 (2014). 10.1016/b978-0-12-420118-7.00008-1

47 Cohen, D. Clozapine and Gastrointestinal Hypomotility. CNS Drugs 31, 1083–1091 (2017). 10.1007/s40263-017-0481-5

48 Seifi, M. et al. Molecular and functional diversity of GABA-A receptors in the enteric nervous system of the mouse colon. J Neurosci 34, 10361–10378 (2014). 10.1523/jneurosci.0441-14.2014

49 Liu, S. et al. Distribution and chemical coding of corticotropin-releasing factor-immunoreactive neurons in the guinea pig enteric nervous system. J Comp Neurol 494, 63–74 (2006). 10.1002/cne.20781

50 Wood, J. D. Enteric Nervous System: Neuropathic Gastrointestinal Motility. Digestive Diseases and Sciences 61, 1803–1816 (2016). 10.1007/s10620-016-4183-5

51 Vance, M. L. Growth-hormone-releasing hormone. Clin Chem 36, 415–420 (1990).

52 Singh, A., Dawson, T. M. & Kulkarni, S. Neurodegenerative disorders and gut-brain interactions. J Clin Invest 131 (2021). 10.1172/JCI143775

53 Challis, C. et al. Gut-seeded alpha-synuclein fibrils promote gut dysfunction and brain pathology specifically in aged mice. Nat Neurosci 23, 327–336 (2020). 10.1038/s41593-020-0589-7

54 Corbille, A. G. et al. Biochemical analysis of alpha-synuclein extracted from control and Parkinson’s disease colonic biopsies. Neurosci Lett 641, 81–86 (2017). 10.1016/j.neulet.2017.01.050

55 Kim, S. et al. Transneuronal Propagation of Pathologic alpha-Synuclein from the Gut to the Brain Models Parkinson’s Disease. Neuron 103, 627–641 e627 (2019). 10.1016/j.neuron.2019.05.035

56 Konings, B. et al. Gastrointestinal syndromes preceding a diagnosis of Parkinson’s disease: testing Braak’s hypothesis using a nationwide database for comparison with Alzheimer’s disease and cerebrovascular diseases. Gut, gutjnl-2023-2329 (2023). 10.1136/gutjnl-2023-329685

57 Ramalingam, N. & Dettmer, U. α-Synuclein serine129 phosphorylation – the physiology of pathology. Molecular Neurodegeneration 18, 84 (2023). 10.1186/s13024-023-00680-x

58 Barrenschee, M. et al. Distinct pattern of enteric phospho-alpha-synuclein aggregates and gene expression profiles in patients with Parkinson’s disease. Acta Neuropathologica Communications 5, 1 (2017). 10.1186/s40478-016-0408-2

59 Tran, D. H. et al. De novo and salvage purine synthesis pathways across tissues and tumors. Cell 187, 3602–3618.e3620 (2024). 10.1016/j.cell.2024.05.011

60 Pareek, V., Pedley, A. M. & Benkovic, S. J. Human de novo purine biosynthesis. Crit Rev Biochem Mol Biol 56, 1–16 (2021). 10.1080/10409238.2020.1832438

61 Mars, R. A. T. et al. Longitudinal Multi-omics Reveals Subset-Specific Mechanisms Underlying Irritable Bowel Syndrome. Cell 182, 1460–1473.e1417 (2020). 10.1016/j.cell.2020.08.007

62 Dewulf, J. P., Marie, S. & Nassogne, M.-C. Disorders of purine biosynthesis metabolism. Molecular Genetics and Metabolism 136, 190–198 (2022). 10.1016/j.ymgme.2021.12.016

63 Smith, W. & Desprez, C. Symptoms of constipation in autistic adults: A systematic literature review on diagnostic methods and presence of actual symptoms. Autism, 13623613241289114 (2024). 10.1177/13623613241289114

64 Cingolani, F., Balasubramaniam, A. & Srinivasan, S. Molecular mechanisms of enteric neuropathies in high-fat diet feeding and diabetes. Neurogastroenterol Motil, e14897 (2024). 10.1111/nmo.14897

65 Finsterer, J. & Strobl, W. Gastrointestinal involvement in neuromuscular disorders. J Gastroenterol Hepatol 39, 1982–1993 (2024). 10.1111/jgh.16650

66 Nezami, B. G. et al. MicroRNA 375 mediates palmitate-induced enteric neuronal damage and high-fat diet-induced delayed intestinal transit in mice. Gastroenterology 146, 473–483 e473 (2014). 10.1053/j.gastro.2013.10.053

67 Boschetti, E. et al. Evidence of enteric angiopathy and neuromuscular hypoxia in patients with mitochondrial neurogastrointestinal encephalomyopathy. Am J Physiol Gastrointest Liver Physiol 320, G768–G779 (2021). 10.1152/ajpgi.00047.2021

68 Zhang, L. et al. Lactate transported by MCT1 plays an active role in promoting mitochondrial biogenesis and enhancing TCA flux in skeletal muscle. Science Advances 10, eadn4508 (2024). doi:10.1126/sciadv.adn4508

69 Walters, D. K., Arendt, B. K. & Jelinek, D. F. CD147 regulates the expression of MCT1 and lactate export in multiple myeloma cells. Cell Cycle 12, 3175–3183 (2013). 10.4161/cc.26193

70 Doherty, J. R. & Cleveland, J. L. Targeting lactate metabolism for cancer therapeutics. The Journal of Clinical Investigation 123, 3685–3692 (2013). 10.1172/JCI69741

71 Xu, Q., Zhu, W. & Yin, Y. Diagnostic value of anti-mitochondrial antibody in patients with primary biliary cholangitis: A systemic review and meta-analysis. Medicine (Baltimore) 102, e36039 (2023). 10.1097/md.0000000000036039

72 Wilschanski, M. et al. A novel familial mutation in the PCSK1 gene that alters the oxyanion hole residue of proprotein convertase 1/3 and impairs its enzymatic activity. PLoS One 9, e108878 (2014). 10.1371/journal.pone.0108878

73 Weaver, A. M. & Patton, J. G. Argonautes in Extracellular Vesicles: Artifact or Selected Cargo? Cancer Res 80, 379–381 (2020). 10.1158/0008-5472.Can-19-2782

74 Parathan, P., Wang, Y., Leembruggen, A. J., Bornstein, J. C. & Foong, J. P. The enteric nervous system undergoes significant chemical and synaptic maturation during adolescence in mice. Dev Biol 458, 75–87 (2020). 10.1016/j.ydbio.2019.10.011

75 Hotta, R. et al. Transplanted progenitors generate functional enteric neurons in the postnatal colon. J Clin Invest 123, 1182–1191 (2013). 10.1172/JCI65963

76 Jiang, M. et al. Calretinin, S100 and protein gene product 9.5 immunostaining of rectal suction biopsies in the diagnosis of Hirschsprung’ disease. Am J Transl Res 8, 3159–3168 (2016).

77 Geleff, S., Schoppmann, S. F. & Oberhuber, G. Increase in podoplanin-expressing intestinal lymphatic vessels in inflammatory bowel disease. Virchows Archiv 442, 231–237 (2003). 10.1007/s00428-002-0744-4

78 Bonfiglio, F. et al. GWAS of stool frequency provides insights into gastrointestinal motility and irritable bowel syndrome. Cell Genomics 1 (2021). 10.1016/j.xgen.2021.100069

79 Boesmans, W., Gomes, P., Janssens, J., Tack, J. & Vanden Berghe, P. Brain-derived neurotrophic factor amplifies neurotransmitter responses and promotes synaptic communication in the enteric nervous system. Gut 57, 314–322 (2008). 10.1136/gut.2007.131839

80 Chai, N. L. et al. Effects of neurotrophins on gastrointestinal myoelectric activities of rats. World J Gastroenterol 9, 1874–1877 (2003). 10.3748/wjg.v9.i8.1874

81 Coulie, B. et al. Recombinant human neurotrophic factors accelerate colonic transit and relieve constipation in humans. Gastroenterology 119, 41–50 (2000). 10.1053/gast.2000.8553

82 Huang, M. et al. Neuronal SNAP-23 is critical for synaptic plasticity and spatial memory independently of NMDA receptor regulation. iScience 26, 106664 (2023). 10.1016/j.isci.2023.106664

83 Suh, Y. H. et al. A neuronal role for SNAP-23 in postsynaptic glutamate receptor trafficking. Nature Neuroscience 13, 338–343 (2010). 10.1038/nn.2488

84 Dolai, S. et al. Pancreas-specific SNAP23 depletion prevents pancreatitis by attenuating pathological basolateral exocytosis and formation of trypsin-activating autolysosomes. Autophagy 17, 3068–3081 (2021). 10.1080/15548627.2020.1852725

85 Tang, Q. et al. Alpha-Synuclein defects autophagy by impairing SNAP29-mediated autophagosome-lysosome fusion. Cell Death & Disease 12, 854 (2021). 10.1038/s41419-021-04138-0

86 Kuo, Y. M. et al. Extensive enteric nervous system abnormalities in mice transgenic for artificial chromosomes containing Parkinson disease-associated alpha-synuclein gene mutations precede central nervous system changes. Hum Mol Genet 19, 1633–1650 (2010). 10.1093/hmg/ddq038

87 Latchman, D. S. Cardiotrophin-1 (CT-1): a novel hypertrophic and cardioprotective agent. Int J Exp Pathol 80, 189–196 (1999). 10.1046/j.1365-2613.1999.00114.x

88 Sánchez-Garrido, A. I. et al. Preventive Effect of Cardiotrophin-1 Administration before DSS-Induced Ulcerative Colitis in Mice. J Clin Med 8 (2019). 10.3390/jcm8122086

89 Kulkarni, S., Kurapati, S. & Bogunovic, M. ’Nervous’ Immunity: Walking the Tightrope. Trends Immunol 41, 359–362 (2020). 10.1016/j.it.2020.03.009

90 Graham, K. D. et al. Robust, 3-Dimensional Visualization of Human Colon Enteric Nervous System Without Tissue Sectioning. Gastroenterology 158, 2221–2235.e2225 (2020). 10.1053/j.gastro.2020.02.035

91 Dershowitz, L. B., Li, L., Pasca, A. M. & Kaltschmidt, J. A. Anatomical and functional maturation of the mid-gestation human enteric nervous system. Nat Commun 14, 2680 (2023). 10.1038/s41467-023-38293-z

92 Kiemen, A. L. et al. CODA: quantitative 3D reconstruction of large tissues at cellular resolution. Nat Methods 19, 1490–1499 (2022). 10.1038/s41592-022-01650-9

